# Microbial metabolite deoxycholic acid controls *Clostridium perfringens*-induced chicken necrotic enteritis through attenuating cyclooxygenase signaling

**DOI:** 10.1101/416107

**Authors:** Hong Wang, Juan D. Latorre, Mohit Bansal, Mussie Abraha, Bilal Al-Rubaye, Guillermo Tellez, Billy Hargis, Xiaolun Sun

**Author notes:** Correspondence: To whom correspondence should be addressed: Center of Excellence for Poultry Science, 1260 W Maple St. O-409, Fayetteville, AR 72701.

## Abstract

*Clostridium perfringens-*induced necrotic enteritis (NE) has reemerged as a prevalent chicken disease worldwide due to reduced usage of prophylactic antibiotics. The lack of antimicrobial alternative strategies to control NE is mainly due to limited insight into the disease pathogenesis. The aim of this study is to investigate the role of microbiota metabolic product secondary bile acid deoxycholic acid (DCA) on preventing NE. *C. perfringens* growth was inhibited by 82.8% in 50 μM DCA Tryptic Soy Broth. Sequential *Eimeria maxima* and *C. perfringens* challenges induced acute NE showed as severe intestinal inflammation and body weight (BW) loss in broiler chickens, while 1.5 g/kg DCA diet dramatically reduced the disease. At the cellular level, DCA alleviated NE-associated ileal epithelial death and reduced lamina propria cell apoptosis. Interestingly, DCA reduced *C. perfringens* invasion into ileum without altering the bacterial ileal luminal colonization. Molecular analysis showed that DCA reduced inflammatory mediators of *Infγ*, *Litaf*, and *Mmp9* mRNA accumulation in ileal tissue. Mechanism studies revealed that *C. perfringens* induced elevated expression of inflammatory mediators of *Infγ*, *Litaf*, *Mmp9,* and *Ptgs2* (Cyclooxygenase- 2 (COX-2) gene) in chicken splenocytes. Blocking COX signaling by pharmacological inhibitor aspirin attenuated INFγ-induced inflammatory response in the splenocytes. Consistent with the *in vitro* assay, chickens fed 0.12 g/kg aspirin diet protected the birds against NE-induced ileal inflammation, intestinal cell apoptosis, and BW loss. In conclusion, microbial metabolic product DCA prevents NE-induced ileal inflammation and BW loss through attenuating inflammatory response. These novel findings offer new strategies against *C. perfringens*-induced diseases.

**Significance Statement:** Widespread antimicrobial resistance has become a serious challenge to both agricultural and healthcare industries. Withdrawing antimicrobials without effective alternatives exacerbates chicken productivity loss at billions of dollars every year, caused by intestinal diseases, such as coccidiosis-and *C. perfringens*-induced necrotic enteritis. This study revealed that microbial metabolic product secondary bile acid DCA prevents *C. perfringens*-induced intestinal disease in chickens through modulating inflammatory COX signaling pathways. Therefore, microbiome and its downstream targets of host inflammatory responses could be used to control NE. These findings have opened new avenues for developing novel antimicrobial free alternatives to prevent or treat *C. perfringens*-induced diseases.

## Background

Antimicrobial resistance is one of the emerging challenges requiring immediate and sustainable counter-actions from agriculture to healthcare (1). Increasing antimicrobial resistance has caused the emergence of multiple drug-resistant microbes or “superbugs”. Recently, a “superbug” of an *Escherichia coli* strain resistant to the last resort antibiotic, Colistin, was reported in USA (2). Overuse of antimicrobial agents in medical and agricultural practice is contributing to exacerbating the episodes of emerging antimicrobial resistant microbes (1). Withdrawing antimicrobials in chicken production, however, has caused new problems for the chicken industry by reducing production efficiency and increasing diseases, such as *Eimeria maxima*- and *Clostridium perfringens*-induced necrotic enteritis (NE) (3). NE is a multi-factorial disease and has a significant economic impact to the chicken industry with annual loss at billions of dollars (4). Subclinical NE (cholangiohepatitis) incidences doubled from 2013 to 2014 in south-eastern Norway were associated with reduced in-feed antimicrobial usage (5), although no major worldwide clinical NE outbreak has recently been reported. Clinical signs of acute NE include watery to bloody (dark) diarrhea, severe depression, decreased appetite, closed eyes, and ruffled feathers. Dissection of dead or severely ill birds shows that the intestine is often distended with gas, very friable, and contains a foul-smelling brown fluid, with clearly visible necrotic lesions (6). At the cellular level, NE birds display intestinal inflammation with diffuse and coagulative necrosis of villi and crypts, infiltration of immune cells into lamina propria, and crypt abscesses (7, 8). Although progress has been made toward understanding risk factors influencing the outcome of NE such as *C. perfringens* virulence, coccidiosis, and feed (9), few effective non-antimicrobial strategies are available.

The human and animal intestine harbors up to trillions of microbes and this intestinal microbiota regulates various host functions such as the intestinal barrier, nutrition and immune homeostasis (10-12). The enteric microbiota regulates granulocytosis and neonatal response to *Escherichia coli K1* and *Klebsiella pneumoniae* sepsis (13), suggesting the key role of the microbiota in protecting the host against systemic infection. At the gut level, fecal transplantation was reported decades ago to prevent *Salmonella infantis* chicken infection (14). More recently, microbiota transplantation has shown tremendous success against recurrent human *Clostridium difficile* infection (15) and *Clostridium scindens* metabolizing secondary bile acids have been shown to inhibit *C. difficile* infection (16). Bile acids synthesized in the liver are released in the intestine and metabolized by gut microbiota into final forms of secondary bile acids (17). Bile acids, particularly the secondary bile acid DCA, are associated with a variety of human chronic diseases, such as obesity, diabetes, and colorectal tumorigenesis (18, 19). Recently, mouse anaerobes and their metabolic product DCA has been found to prevent and treat *Campylobacter jejuni*-induced intestinal inflammation in germ-free mice through attenuating host inflammatory signaling pathways (20). However, what is the role of bile acids in NE is unknown.

Coccidiosis is associated with strong immune response and intestinal inflammation in chickens (21, 22). *C. perfringens* produces various toxins (23) and induces hemolysis, epithelial barrier dysfunction, tissue necrosis and severe inflammation in non-chicken models (24, 25). Among inflammatory signaling pathways, cyclooxygenases (COX)-catalyzed prostanoids regulate various activities including cell proliferation, apoptosis and migration (26), gastrointestinal secretion (27), body temperature (28), inflammation (29), and pain sensation (30). COX-1 and COX-3 (gene *Ptgs1*, alternative splicing) constitutively expressed are important for intestinal integrity. Inducible COX-2 (*Ptgs2*) activity is associated with various inflammatory diseases including inflammatory bowel disease (31) and radiation-induced small bowel injury (32). COX-2 increases gut barrier permeability and bacterial translocation across the intestinal barrier (33, 34). Paradoxically, COX-2 enhances inflammation resolution through prostaglandin D2 (35). Although non-selective COX inhibitor aspirin is used to prevent various chronic diseases, it inflicts intestinal inflammation to the healthy intestine (36). However, the role of COX signaling on NE remains elusive.

Currently, limited knowledge is available on the relationship between NE pathogenesis, the microbiome, and host inflammatory response. Because DCA attenuates *C. jejuni*-induced intestinal inflammation (20) and prevents *in vitro* growth of *C. difficile* (16), a same genus member of *C. perfringens*, we hypothesized that the microbiota metabolic product DCA attenuated NE through inhibiting intestinal inflammation. We found that DCA decreased NE-induced intestinal inflammation, *C. perfringens* invasion, intestinal cell death, and body weight loss. Blocking the inflammatory COX signaling pathways by aspirin reduced NE-induced intestinal inflammation. These novel findings of microbiome DCA and COX inhibitor against NE offer new strategies to prevent and treat *C. perfringens*-induced diseases.

## Results

### DCA prevents *C. perfringens in vitro* growth

Based on previous findings and current state of knowledge, we reasoned that DCA would prevent *C. perfringens* growth. To test this hypothesis, *in vitro* inhibition experiments were conducted, in which *C. perfringens* was inoculated in Tryptic Soy Broth (TSB) with sodium thioglycollate under anaerobic condition. The TSB was also added with various concentrations of bile acids, including conjugated primary bile acid taurocholic acid (TCA), primary bile acid cholic acid (CA), and secondary bile acid DCA. The results showed that DCA inhibited *C. perfringens* growth at 0.01 (−33.8%) and 0.05 mM (−82.8%, clear broth), respectively, compared to control, while TCA (−16.4%) and CA (−8.2%) didn’t prevent the bacterial growth (cloudy broth) even at 0.2 mM (Figure 1A and B). We then examined whether other secondary bile acids were also bacteriostatic in TSB. Interestingly, *C. perfringens* growth inhibited by lithocholic acid (LCA; - 22.6 and -23.8%) and ursodeoxycholic acid (UDCA; -10.0 and -25.3%) at 0.2 and 1 mM, respectively (Figure 1C and D) was far less effective compared to DCA. These results suggest that the secondary bile acid DCA is more efficient to prevent *C. perfringens in vitro* growth compared to other bile acids.

**Figure 1.**
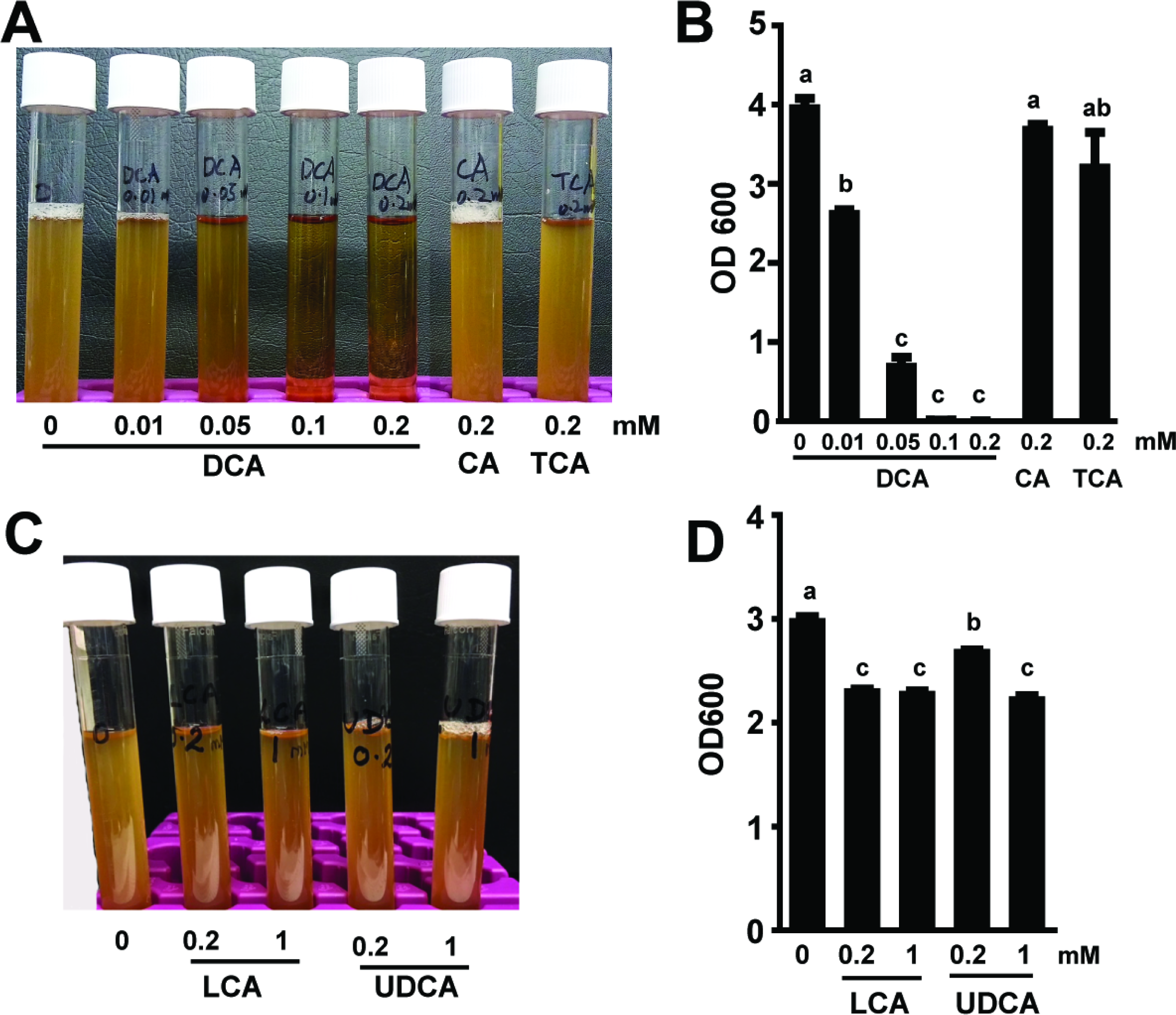
DCA inhibits *C. perfringens in vitro* growth. *C. perfringens* (10^3^ CFU) were inoculated into TSB supplemented with various concentrations of conjugated primary bile acid TCA, primary bile acid CA, and secondary bile acids DCA, LCA, and UDCA. (A) Image of bile acids did (clear broth) or didn’t (cloudy broth) inhibit *C. perfringens* growth. (B) OD600 reading of the broth in panel A. (C) Image of other secondary bile acids inhibited *C. perfringens* growth at less efficiency compared to DCA. (B) OD600 reading of the broth in panel C. All graphs depict mean ± SEM. Different letters of a, b and c mean P<0.05. Results are representative of 3 independent experiments.

### DCA prevents NE-induced intestinal inflammation in chicken ileum

To further address whether DCA reduced coccidia *E. maxima*- and *C. perfringens*-induced NE in birds, one-day-old broiler chicks were fed with 1.5 g/kg CA or DCA diets and body weight (BW) gain were individually measured. To induce NE, the birds were infected with 20,000 sporulated oocysts/bird *E. maxima* at 18 days of age and then infected with 10^9^ CFU/bird *C. perfringens* at 23 and 24 days of age. Because coccidiosis and NE induce severe intestinal inflammation (7, 8), the impact of DCA on chicken NE was investigated. Upper ileum tissue was collected as Swiss Rolls, processed with H&E staining, and performed histopathology analysis. Notably, *E. maxima* (Em) infection induced severe intestinal inflammation (ileitis) as seen by immune cell infiltration (yellow arrows) into lamina propria, crypt hyperplasia, and mild villus height shortening compared to uninfected birds (Figure 2A). NE birds displayed worse ileitis as seen by necrosis and fusion of villi and crypt (Green arrow), massive immune cell infiltration, and severe villus shortening. In contrast, DCA diet dramatically attenuated NE-induced ileitis and histopathology score (Figure 2B), while CA reduced NE-induced ileitis and histopathology score. These results indicate that DCA protects against coccidiosis- and NE-induced severe ileitis.

**Figure 2.**
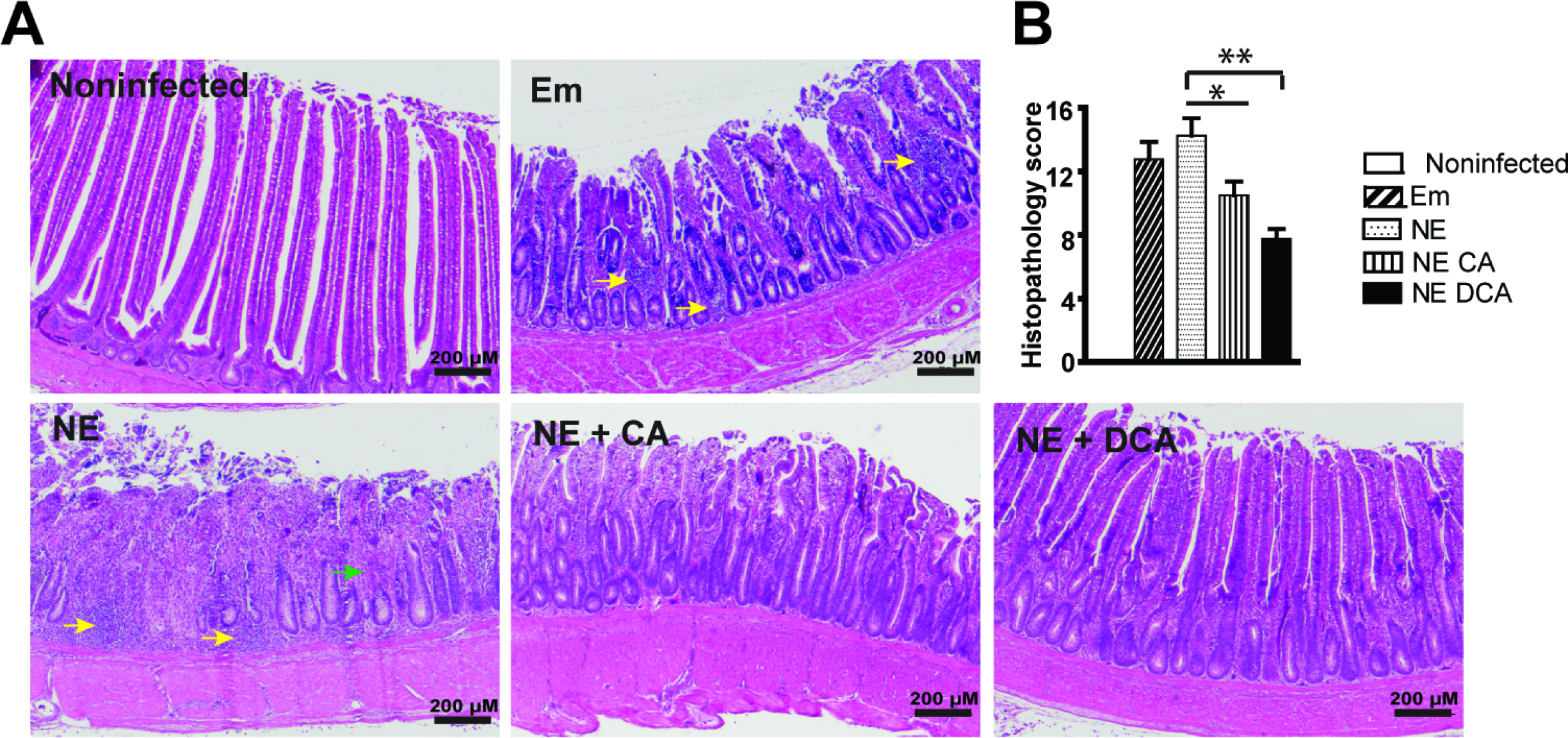
DCA attenuates NE-induced intestinal inflammation. Cohorts of thirteen one-day-old broiler chicks were fed basal, 1.5 g/kg CA or DCA diets. The birds were infected with *E. maxima* at 18 days of age and *C. perfringens* at 23 and 24 days of age. Four birds per group were sacrificed and sampled at 26 days of age. (A) H&E staining showing representative intestinal histology images. (B) Quantification of histological intestinal damage score. Scale bar is 200 μm. All graphs depict mean ± SEM. *, P<0.05; **, P<0.01. Yellow arrows: immune cell infiltration; green arrow: fusion of villi and crypts. Results are representative of 3 independent experiments.

### DCA attenuates NE-induced intestinal cell necrosis and apoptosis

Healthy intestinal epithelial cells have polarity (37) and their nuclei are located toward the basal membrane (38), while stressed dying (apoptosis or necrosis) cells lose polarity (39) and their nuclei disperse from basal to apical membranes (40). It was then sought to examine whether cell death was relevant in DCA attenuating NE-induced ileitis. Since it was difficult to find reliable chicken antibodies to detect apoptosis or necrosis in chicken histology slides, we first resorted to classical histological analysis under high magnification. Consistently, the epithelial nuclei (dark blue) in heathy control bird villi were distributed close to the basal membrane (at the right side of the yellow dash line, Figure 3A lower panel left). In contrast, the nuclei in inflamed villi epithelial cells of Em and NE birds were scattered from basal to the apical membranes, indicating epithelial cell death in villi of those birds. As a contrast, the DCA diet prevented epithelial cell nucleus translocation to apical side, suggesting cell death reduction. To further characterize the villus cell death, terminal deoxynucleotidyl transferase (TdT) dUTP Nick-End Labeling (TUNNEL) assay was used, which detects later stage of cell apoptosis. Consistent with histopathology results, coccidiosis and NE induced scattered (Em birds) or concentrated (NE birds) apoptosis cells (green dots) in villus lamina propria, while cellular apoptosis was attenuated in the DCA treatment birds (Figure 3B). To quantify the apoptosis, ImageJ was used to count apoptosis cells (TUNEL, green) and total cells (DAPI, blue). Consistent with histopathology results, DCA reduced NE-induced cell apoptosis (Figure 3C). These results indicate that DCA prevents against coccidiosis- and NE-induced cell death in intestinal epithelial and lamina propria cells.

**Figure 3.**
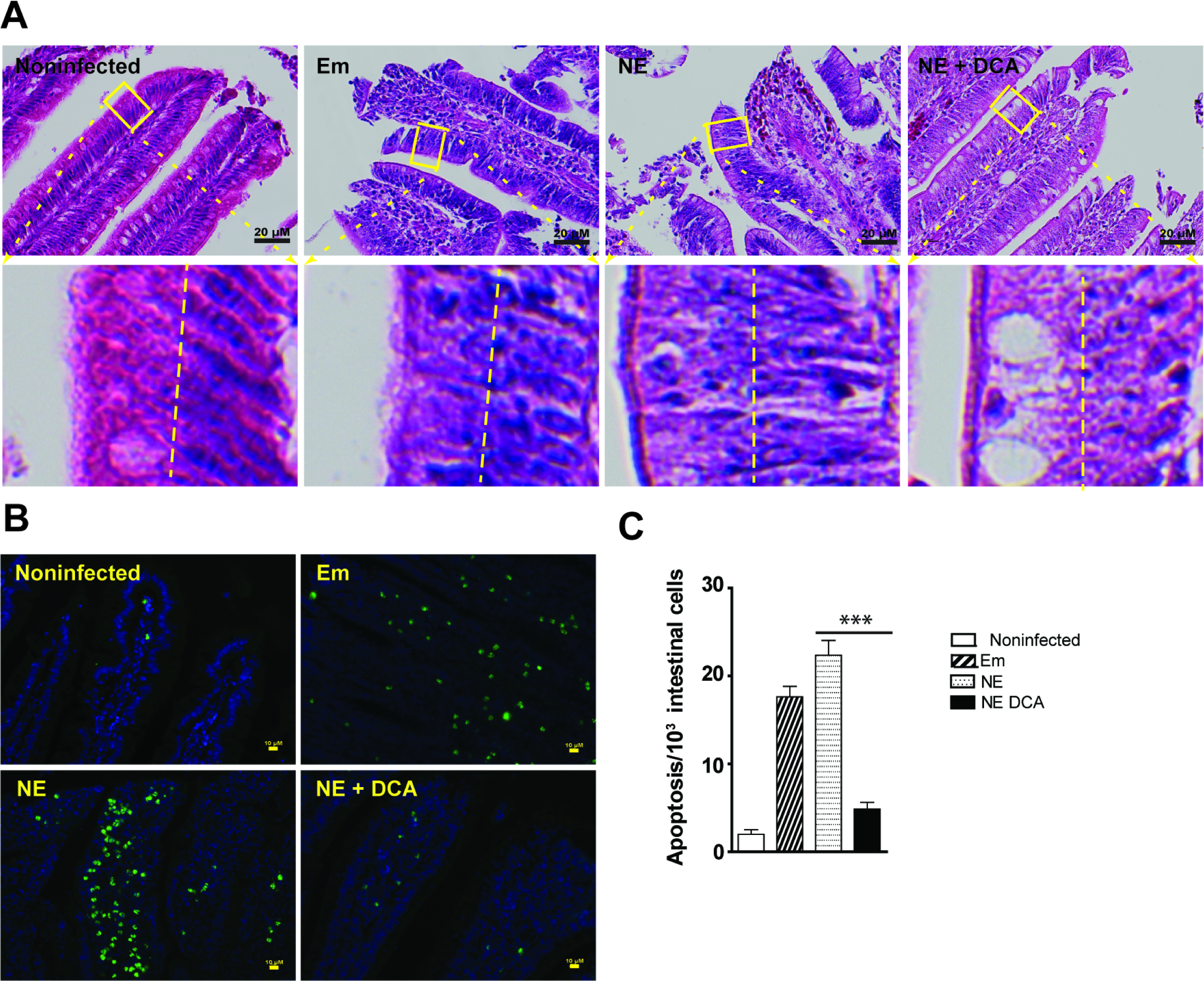
DCA attenuates NE-induced intestinal cell death and apoptosis. Cohorts of broiler chicks were fed different diets, infected, and sampled as in Figure 2. (A) Representative intestinal cell death (deviated nuclei) using H&E staining. (B) Representative villus cell apoptosis (green) using TUNEL assay. (C) Quantified apoptotic cells using ImageJ. Scale bars are 20 μm (A) and 10 μm (B). All graphs depict mean ± SEM. ***, P<0.001. Results are representative of 3 independent experiments.

### DCA reduces *C. perfringens* invasion, NE-induced inflammatory response, and productivity loss

Given DCA inhibited *C. perfringens* growth *in vitro*, it was logic to reason that DCA might also reduce *C. perfringens* intestinal overgrowth in NE birds. To examine this possibility, *C. perfringens* colonization level was measured in the intestinal lumen using real-time PCR of *C. perfringens* 16S rDNA. Surprisingly, ileal luminal *C. perfringens* colonization in DCA birds was not significantly different from NE control birds (Figure 4A), while bird histopathology were distinct between the two groups of birds. We then reasoned that the pathogen invasion into tissue was the main driving factors of NE pathogenesis, but not the pathogen luminal colonization level. To quantify the bacterial invasion, total DNA in ileal tissue was isolated and *C. perfringens* was measured using real time PCR. DCA attenuated more than 95 % of *C. perfringens* invasion into ileal tissue (Figure 4B). To have a better overview of the bacterial tissue invading distribution, we used a fluorescence *in situ* hybridization (FISH) technique. We found that while *C. perfringens* was present deeply in the inflamed villus and crypt lamina propria of NE control birds, the bacterium was barely detectable in the ileal tissue of DCA birds (Figure 4C). Because DCA reduced *C. perfringens* invasion and intestinal inflammation, the impact of DCA on various proinflammatory mediators was evaluated in ileal tissue using Real-Time PCR. *C. perfringens* induced inflammatory *Infγ*, *Litaf* (*Tnfα*), and *Mmp9* mRNA accumulation in chicken ileal tissue, an effect attenuated by 51, 82, and 93%, respectively, in DCA fed chickens (Figure 4D).

**Figure 4.**
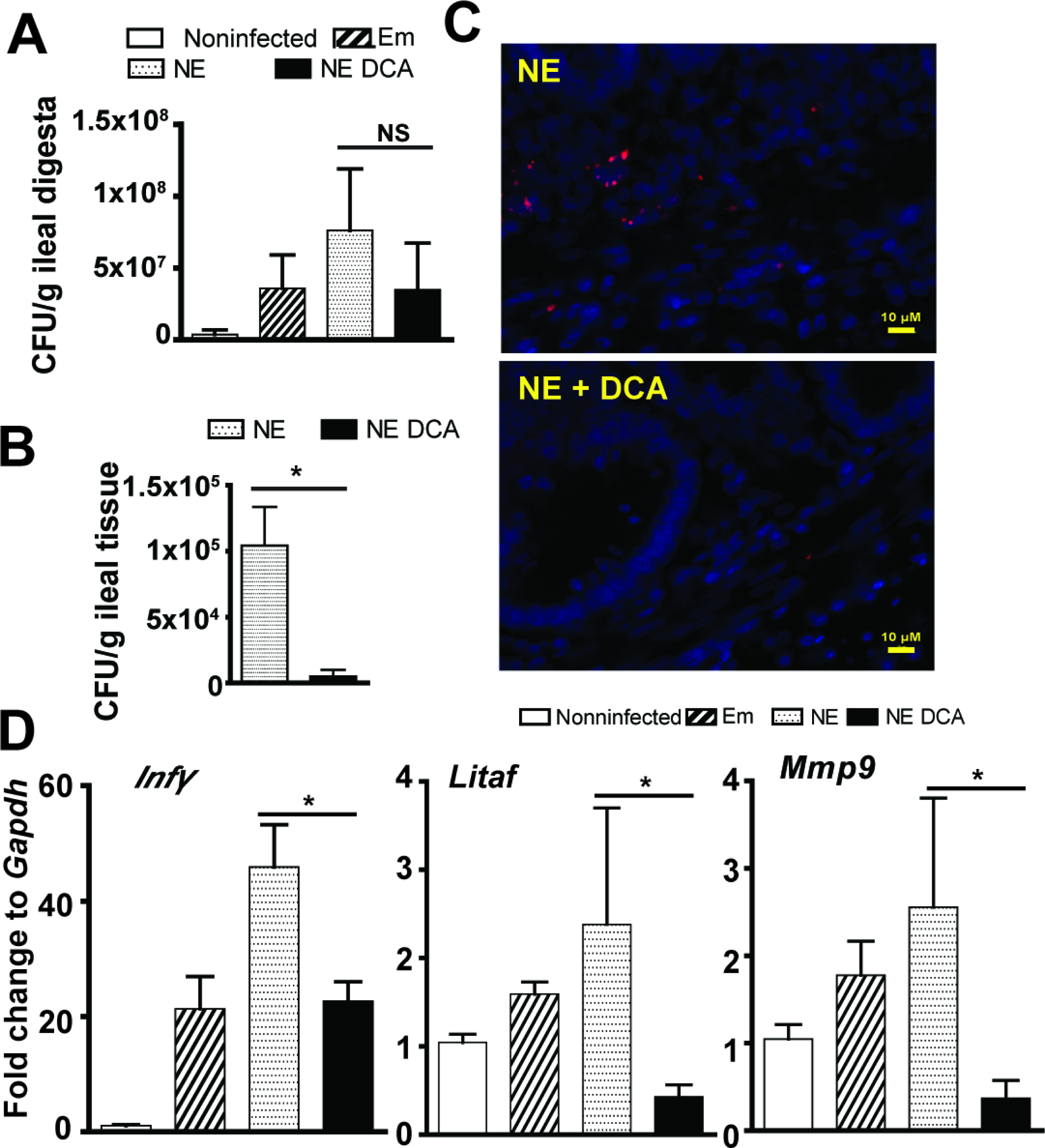
DCA reduces *C. perfringens* invasion and inflammatory response. Chickens were fed different diets, infected, and sampled as in Figure 2. (A) Luminal *C. perfringens* colonization level quantified by 16S rDNA real-time PCR. (B) *C. perfringens* invasion into intestinal tissue quantified by 16S RNA real-time PCR. (C) Presence of *C. perfringens* (red dots) in ileal sections of NE and NE+DCA birds, detected using fluorescence in situ hybridization (FISH) assay. (D) Ileal *Infγ*, *Litaf* (*Tnfα*), and *Mmp9* mRNA qPCR fold change relative to uninfected birds and normalized to *Gapdh*. Scale bar is 10 μm. All graphs depict mean ± SEM. NS, not significant; *, P<0.05. Results are representative of 3 independent experiments.

Body weight (BW) is one of important productivity parameters for meat chickens and reflects collective response during NE. The productivity results showed that DCA (solid black bar) but not CA (vertical-line bar) diet promoted bird daily BW gain during 0-18 days of age compared to birds fed control diets (open, tilted line, and dotted bars, Supplemental Figure 1). BW gain was reduced in birds infected with *E. maxima* (Em) (tilted line and dotted bars) during 18-23 days of age (coccidiosis phase) compared to noninfected birds (open bar). Subsequent *C. perfringens* infection reduced NE control birds (dotted bar) BW gain during 23-26 days of age (NE phase) compared to Em birds (tilted line bar). Consistent with histopathology analysis, DCA prevented productivity loss at coccidiosis and NE phases compared to the NE control birds. Interestingly, the primary bile acid CA diet attenuated body weight loss at NE phase but failed at coccidiosis phase compared to the NE control birds.

### COX inhibitor aspirin alleviates *C. perfringens*-induced inflammatory response in splenocytes

Inflammatory events shape intestinal diseases and targeting the inflammatory response attenuates disease progress such as in Inflammatory Bowel Disease (41) and campylobacteriosis (42). To dissect how the host inflammatory response is involved in NE-induced ileitis, a primary chicken splenocyte cell culture system was then used similarly to previous report (43). After isolation from chickens, the splenocytes were infected with *C. perfringens* (MOI 100) for 4 hours. The results showed that *C. perfringens* increased inflammatory mediators of *Infγ*, *Litaf* (*Tnfα*), *Mmp9* and *Ptgs2* (protein COX-2) mRNA accumulation by 1.54, 1.69, 1.72 and 8.65 folds, respectively, compared to uninfected splenocytes (Figure 5A). Because COX-2 are important mediators in the inflammatory response (31), COX inhibitor aspirin was then used in *C. perfringens*-infected chicken splenocytes. Interestingly, aspirin failed to reduce *C. perfringens*-induced inflammatory gene expression (data not shown). We then reasoned that COX signaling acted on *C. perfringens*-induced inflammatory cytokines. Inflammatory cytokine of recombinant chicken INFγ (ch-INFγ) was then used to challenge splenocytes in the presence of aspirin. Aspirin reduced ch-INFγ-induced inflammatory gene expression of *Infγ*, *Litaf* (*Tnfα*), and *Mmp9* by 44, 45, and 65%, respectively (Figure 5B). Because no chicken TNFα was available, murine INFγ (mINFγ) and mTNFα were used. Consistently, aspirin also reduced murine INFγ (mINFγ)-induced inflammatory gene expression of *Infγ*, *Litaf*, and *Mmp9* by 41, 27, and 45%, respectively (Supplemental Figure 2A). Similarly, aspirin reduced mTNFα-induced inflammatory gene expression of *Infγ*, *Litaf*, and *Mmp9* by 49, 53, and 27%, respectively (Supplemental Figure 2B). These data indicate that *C. perfringens* induces inflammatory cytokines and COX-2 while blocking COX signaling by aspirin reduces inflammatory cytokine-induced responses, suggesting that aspirin poses protection potential against NE detrimental inflammatory response.

**Figure 5.**
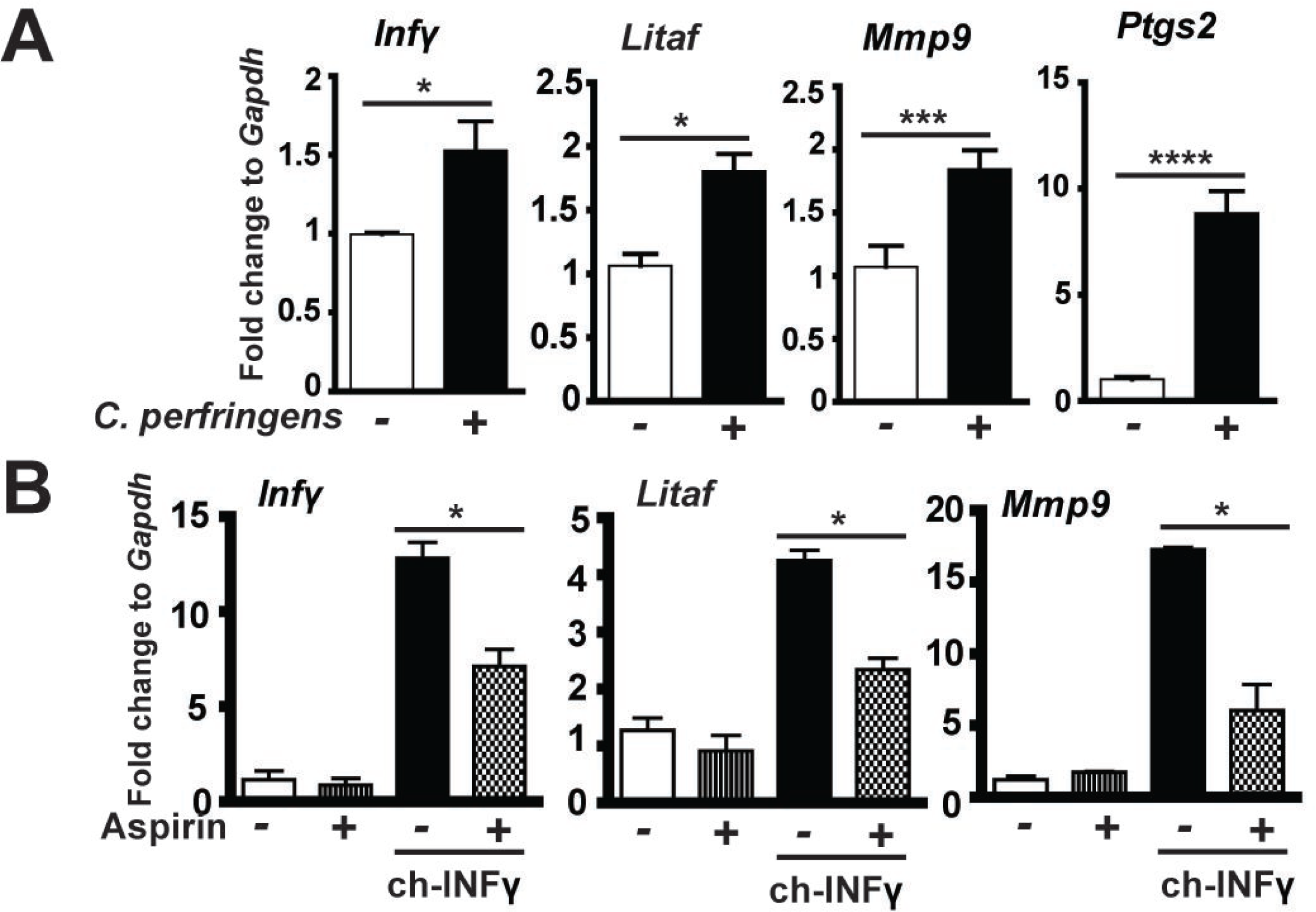
COX inhibitor aspirin alleviates *C. perfringens*-induced inflammatory response in chicken splenocytes. Splenocytes isolated from broiler chickens were infected with *C. perfringens* (MOI 100) for 4 hr or stimulated with chicken INFγ (1μg/ml) for 2 hr in the presence of 1.2 mM aspirin. RNA was extracted, reverse-transcribed, and quantified using a Bio-Rad 384 PCR platform. (A) *Infγ*, *Litaf*, *Mmp9*, and *Ptgs2* mRNA fold change normalized to *Gapdh*. (B) Gene expression fold change in the presence of chicken INFγ (ch-INFγ) and aspirin. All graphs depict mean ± SEM. *, P<0.05; ***, P<0.001. Results are representative of 2 independent experiments.

### Aspirin attenuates NE-induced ileitis, intestinal cell apoptosis, and productivity loss

To functionally assess the protective effect of aspirin against NE-induced ileitis, broiler chickens fed with 0.12 g/kg aspirin diet (ASP) were infected with *E. maxima* and *C. perfringens* as describe before. Ileal tissue were collected and histopathology analysis was performed to assess NE. Consistently, NE birds had severe ileal necrosis with immune cell infiltration and villus shortening (Figures 6A). Notably, ASP attenuated NE-induced intestinal inflammation and histopathological score (Figures 6A and B). At cellular level, ASP reduced NE-induced immune cell apoptosis in villus lamina propria (Figure 6 C and D). On growth performance, ASP birds grew slower compared to control diet birds during 0-18 days of age (Supplemental Figure 3). This is because aspirin inhibits all COX isoforms and COX-1 and -3 are important for intestinal homeostasis and growth. Notably, ASP attenuated NE-induced BW loss by 60% during NE phase of 23-26 days of age, while no difference between ASP and NE birds during coccidiosis phase of 18-23 days of age. These data suggest that aspirin attenuates NE-induced intestinal inflammatory response, intestinal cell death, and BW loss.

**Figure 6.**
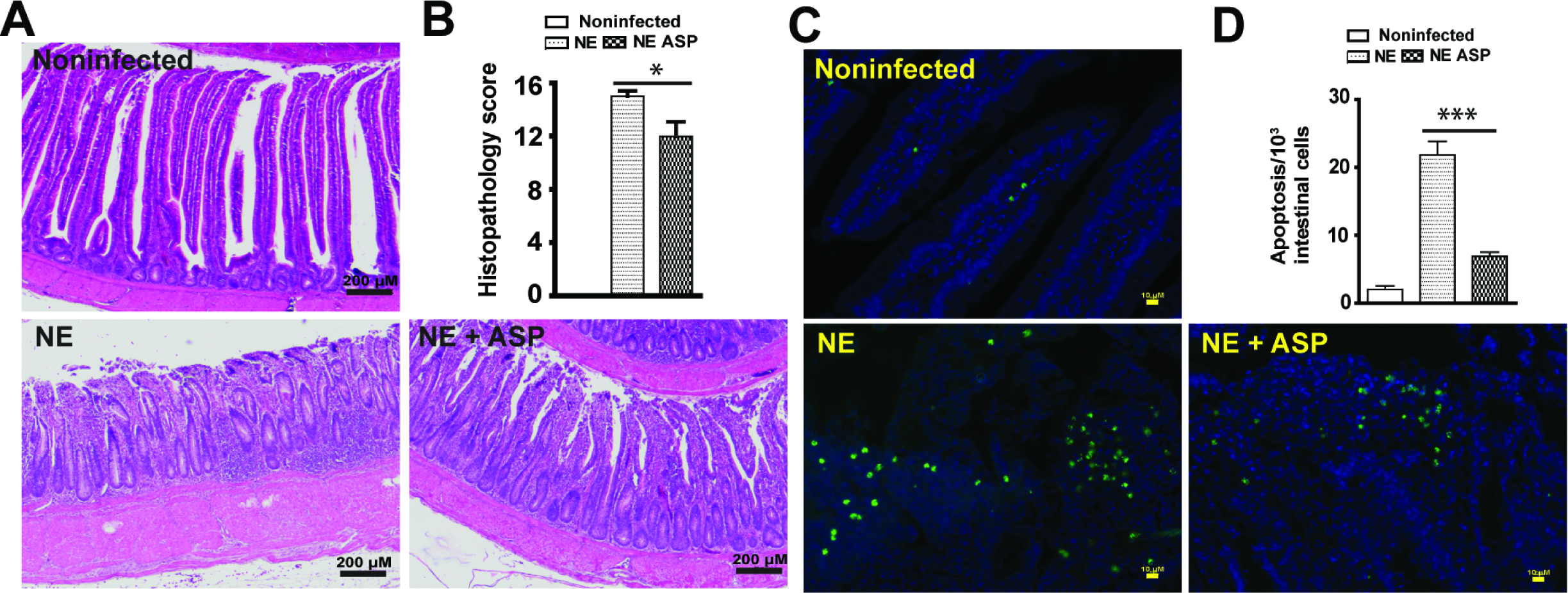
Aspirin attenuates NE-induced intestinal inflammation and apoptosis. Cohorts of thirteen chickens were fed basal and 0.12 g/kg aspirin diet. The birds were infected as in Figure 2 and four birds per group were sampled at 26 days of age. (A) H&E staining showing representative intestinal histology images. (B) Quantification of histological intestinal damage score. (C) Representative cell apoptosis (green) using TUNEL assay. (D) Quantification of apoptotic cells. Scale bars are 200 μm in panel A and 10 μm in panel C. All graphs depict mean ± SEM. *, P<0.05; ***, P<0.001. Results are representative of 2 independent experiments.

## Discussion

Although NE has reemerged as a prevalent chicken disease worldwide in the antimicrobial free era (5), the lack of comprehensive molecular mechanism insight into NE severely hinders the development of antimicrobial alternatives to control this disease (44). Many virulent factors of *Eimeria* and *C. perfringens* are identified but few findings are effective to control NE in chicken production (9, 45), suggesting that important players/factors in NE pathogenesis were overlooked, such as microbiome and host response. We reported here that microbiota metabolic product DCA attenuated chicken NE by reducing chicken inflammatory COX signaling pathways. These new findings offer approaches for exploring novel antimicrobial alternatives to control *C. perfringens*-induced diseases.

It is a relatively new concept to manipulate microbiota and its metabolic products against infectious diseases. Fecal transplantation was used in chickens decades ago to prevent *S. infantis* infection (14). Microbiome plays an important role in susceptibility to human *C. difficile* infection (46). Anaerobe *C. scindens*-transformed secondary bile acids prevent *C. difficile* germination and growth (16). Lithocholic acid and DCA but not primary bile acid CA inhibit *C. difficile* vegetable growth and toxin production (16, 47). However, whether secondary bile acids prevent or treat *C. difficile* infection in human or animal models is still unknown. It has been recently found that orally gavaging DCA attenuated *C. jejuni*-induced intestinal inflammation in germ-free mice (20). Based on the knowledge, we then reasoned that DCA might prevent *E. maxima*-and *C. perfringens*-induced chicken NE. Indeed, dietary DCA prevented NE and its associated productivity loss. The reduction of ileitis was coupled with reduced *C. perfringens* intestinal tissue invasion and intestinal inflammation and cell death. Intriguingly, DCA failed to reduce *C. perfringens* ileal luminal colonization, suggesting that the mechanism of DCA action is independent of intestinal luminal colonization exclusion and is possibly through modulating other factors such as inflammation.

At the cellular level, the intestinal tract of NE-inflicted birds displays severe small intestinal inflammation, showing massive immune cells infiltration into lamina propria, villus breakdown, and crypt hyperplasia (48, 49). Intestinal inflammation is critical to clear invaded microbes and to resolve inflammation, while overzealous inflammation causes more bacterial invasion and further collateral damage and inflammation (50). Infectious bacteria often hijack the inflammatory pathways to gain survival and invasion advantage. For example, *Salmonella* Typhimurium induces extensive inflammation in mouse intestine and thrives on the inflammation (51). Interestingly, *S.* Typhimurium infection causes immunosuppressive effect in neonatal chickens because of lymphocyte depletion (52). Unlike *S.* Typhimurium infection in neonatal birds, coccidia infection induces strong immune response and intestinal inflammation in chickens (21, 22). Furthermore, NE birds suffered more intestinal inflammation compared to Em birds in current study. Consistent with the “over-inflammation” model, NE birds with severe intestinal inflammation showed extensive immune cell infiltration, increased inflammatory cytokine gene expression, and epithelial cell hyperplasia and death. Conversely, DCA attenuated the intestinal inflammation and inflammatory cytokines and improved the growth performance and reduced the NE pathology. Consistent with the reasoning of anti-inflammation reducing NE, blocking inflammatory COX-2 signaling pathway by aspirin alleviated intestinal inflammation, villus apoptosis and NE-induced BW loss. These findings indicate that DCA attenuates NE through decreasing inflammatory signaling pathways.

Different strains of *C. perfringens* produce a variety of toxins including alpha (CPA), beta (CPB), epsilon (ETX), iota (ITX), enterotoxin (CPE), necrotic enteritis B-like toxin (NetB), and others (53). Among them, researchers have reported that NetB but not CPA induced NE in chickens (54). However, NE doesn’t have strong association with NetB positive *C. perfringens* in US (55). In our NE model, no difference of *NetB* gene expression in ileal digesta between healthy control and NE birds (data not shown) suggests limited role of the toxin. Generally, *C. perfringens* toxins induce cell death of apoptosis, necrosis, necroptosis, and pyroptosis in the presence of influxed extracellular calcium (53). Higher dietary calcium increases chicken mortality and reduces growth performance in coccidiosis-induced NE model (56), suggesting the important role of toxin-induced and calcium-mediated cell death in NE. The reduction of intestinal cell death by DCA and aspirin suggests possible toxin involvement in our NE model. Future work on *C. perfringens* toxins, DCA, and inflammatory response will be conducted.

Altogether, these data reveal that the microbial metabolic product secondary bile acid DCA attenuates NE, through reducing NE-induced host inflammatory response. These findings highlight the importance of elucidating the molecular relationship between infectious pathogen, microbiome, and host response. These discovers could be applied to control NE and other intestinal diseases targeting microbiome and host inflammatory response.

**Abbreviations**
NE: necrotic enteritis
BW: body weight
DCA: deoxycholic acid
CA: cholic acid
COX: cyclooxygenases
TSB: tryptic Soy Broth

## Declarations

### Acknowledgements

The authors would like to thank the support from staff at Poultry Health Laboratory and Feed Mill in Department of Poultry Science at University of Arkansas, Fayetteville. We also thank P. L. Matsler on helping our histology slides.

### Funding

This research was supported by Arkansas Biosciences Institute grant to X. Sun.

### Availability of data and materials

Data sharing is not applicable to this article as no datasets were generated or analyzed during the current study.

### Authors’ contributions

H.W. and X.S. designed the experiments and wrote the manuscript with input from co-authors. H.W. and X.S. performed animal and cell experiments and most analysis with the participation from co-authors. All authors read and approved the final manuscript.

### Ethics approval and consent to participate

All animal protocols were approved by the Institutional Animal Care and Use Committee of the University of Arkansas at Fayetteville.

### Consent for publication

Not applicable.

### Competing interests

The authors declare that they have no competing interests.

## Methods

### Chicken experiment

Cohorts of thirteen one-day-old broiler chicks per group were obtained from Cobb-Vantress (Siloam Springs, AR). Chicks were neck-tagged and randomly allocated to floor pens with new pine shavings as litter in an environmentally controlled isolate room suitable for up to biosafety level (BSL) 3 animal experiments. The birds were provided with their respective diet and water *ad libitum*, and temperature was maintained at 34°C for the first 5 d and was then gradually reduced until a temperature of 23°C was achieved at day 26d of age. The birds were fed a corn-soybean meal-based starter diet during 0-10 days of age and a grower diet during 11-26 days of age. The basal diet was formulated as described before (57). Treatment diets were supplemented with 1.5 g/kg CA or DCA or 0.12 g/kg aspirin (all from Alfa Aesar). No antibiotics, coccidiostats or enzymes were added to the feed. *E. maxima* kindly provided by Dr. John Barta was M6 strain, which was a single oocyst-derived isolate at Ontario Veterinary College in 1973. The *E. maxima* were propagated in chickens as describe before (58). The recovered oocysts were sporulated as previously described (59). *C. perfringens* kindly donated from USDA-ARS, College Park, TX (60, 61) was confirmed alpha-toxin positive using a multiplex PCR assay (60, 62). An aliquot of frozen *C. perfringens* was grown in TSB plus sodium thioglycolate overnight for the NE challenge study, and was serially diluted and plated on tryptic soy agar plus sodium thioglycolate for enumerating CFU. In previous experiments, birds infected with this aliquot of *C. perfringens* alone didn’t show any signs of NE and had comparable body weight gain to noninfected birds (data not showed). In current experiments, birds were infected with 20,000 sporulated oocytes/bird *E. maxima* at 18 days of age and 10^9^ CFU/bird *C. perfringens* at 23 and 24 days of age. Chicken body weight and feed intake were individually measured at d 0, 18, 23, and 26 days of age. Bird health status was monitored daily after the pathogen infection. Four birds with average BW of the treatment group were sacrificed at 23 and 26 days of age. Because gross necrotic lesion was often observed in upper ileum in this NE model, the ileal tissue and digesta samples from all sacrificed birds were collected for RNA and DNA analysis. The ileal tissue of ∼8 cm length was also Swiss-rolled for H&E staining and histopathology analysis. Images were acquired using a Nikon TS2 fluorescent microscope. Ileal inflammation was scored blindly using the H&E Swiss roll slides (4 slides (birds)/treatment). Briefly, each Swiss roll slide was divided into 4 areas and scored. Total histopathological scores were then calculated by adding the four area scores. The following score scales used were based on previous ileitis (63, 64) and colitis scoring systems (43): score 0: no inflammation and villi and crypt intact; score 1: small number infiltration cells in laminar propria of villi and crypts or villi minimally shortened; score 2: more extensive infiltration cells in laminar propria of villi and crypts, villi shortened >¼ and edema, or crypt hyperplasia; score 3: pronounced infiltration cells in laminar propria of villi, crypts, submucosa, and muscularis, villi shortened >½ and edema, or crypt hyperplasia and regeneration; and score 4: necrosis, villus diffuse, ulcers, crypt abscesses, or transmural inflammation (may extend to serosa).

### *C. perfringens*-induced inflammatory response using primary splenocytes

Splenocytes were isolated similarly to described previously (42). Brie?y, chicken spleen was resected, homogenized into splenocytes using frosted glass slides, and pooled together in RPMI 1640 medium supplemented with 2% fetal bovine serum, 2mM L-glutamine, 50 μM 2- mercaptoethanol. After lysed the red blood cells, the collected cells were plated at 2 x 10^6^ Cells/well in 6-well plates. The cells were pre-treated with 1.2 mM aspirin for 45 min. Cells were then challenged with murine INFγ (1μg/ml, Pepro Tech), chicken INFγ (1μg/ml, IBI Scientific), or *C. perfringens* (multiplicity of infection 100). The cells were lysed in TRIzol (Invitrogen) for RNA isolation after 2 or 4 hours of cytokines or *C. perfringens* treatment, respectively.

### Real time RT-PCR of mRNA gene expression and *C. perfringens* quantification

Total RNA from ileal tissue or splenocytes was extracted using TRIzol as described before (20, 65). cDNA was prepared using M-MLV (NE Biolab). mRNA levels of proinflammatory genes were determined using SYBR Green PCR Master mix (Bio-Rad) on a Bio-Rad 384-well Real-Time PCR System and normalized to *Gapdh.* To quantify *C. perfringens* intestinal luminal colonization or tissue invasion, ileal digesta or tissue were weighed, bead-beated, and extracted for DNA using phenol-chloroform method as described before (20). *C. perfringens* in the digesta and tissue was quantified using specific *C. perfringens* 16S rDNA primers by the qPCR. The PCR reactions were performed according to the manufacturer’s recommendation. The following gene primers were used:

*Cp16S*_forward: 5’- CAACTTGGGTGCTGCATTCC-3’; *Cp16S* reverse: 5’- GCCTCAGCGTCAGTTACAG-3’; *Mmp9*_forward: 5’-CCAAGATGTGCTCACCAAGA-3’; *Mmp9*_reverse: 5’-CCAATGCCCAACTTCTCAAT-3’; *Litaf _*forward: 5’- AGATGGGAAGGGAATGAACC; *Litaf _*reverse: 5’-GACGTGTCACGATCATCTGG-3’; *Il1β*_forward: 5’-GCATCAAGGGCTACAAGCTC-3’;

*Il1 β*_reverse: 5’-CAGGCGGTAGAAGATGAAGC-3’; *Infγ_*forward: 5’- AGCCGCACATCAAACACATA -3’; *Infγ_*reverse: 5’-TCCTTTTGAAACTCGGAGGA-3’; *Ptgs2*_forward: 5’-ACCAGCATTTCAACCTTTGC-3’; *Ptgs2*_reverse: 5’- CCAGGTTGCTGCTCTACTCC-3’; *Gapdh_*forward: 5’-GACGTGCAGCAGGAACACTA-3’; *Gapdh*_reverse: 5’- CTTGGACTTTGCCAGAGAGG-3’. Gene expression of fold change was calculated using ΔΔCt method (66) and *Gapdh* as internal control.

### TUNNEL assay

Cell apoptosis in intestinal tissue was visualized using TUNNEL assay. Briefly, ileal tissue slides were deparaffinized with xylene bath for 3 times and rehydrated with 100%, 95%, and 70% ethanol. The tissue was then incubated with TUNNEL solution (5 μM Fluorescein-12-dUTP (Enzo Life Sciences), 10 μM dATP, 1 mM pH 7.6 Tris-HCl, 0.1 mM EDTA, 1U TdT enzyme (Promega) at 37° C for 90 min. The slides were counter-stained with DAPI for nucleus visualization. The fluorescent green apoptosis cells were evaluated and imaged using a Nikon TS2 fluorescent microscopy. The green dots in representative 3 areas per slide were counted as apoptosis cells and blue dots of nuclei were counted as total cells using ImageJ (67) particle analysis and its plugin of Nikon ND2 reader. The results were showed as apoptosis cells per 1,000 total intestinal cells.

### Fluorescence in situ hybridization (FISH)

*C. perfringens* at ileal tissue was visualized using FISH assay similarly to previously described (43). Briefly, tissue slides were deparaffinized, hybridized with the FISH probe, washed, stained with DAPI, and imaged using a Nikon TS2 fluorescent Microscope system. The FISH probe sequence of Cp85aa18: 5’-/Cy3/TGGTTGAATGATGATGCC-3’ (68) was used to probe the presence of *C. perfringens* similar to a previous report (65). Briefly, deparaffinized, formalin-fixed 5 μm thick sections were incubated for 15 minutes in lysozyme (300,000 Units/ml lysozyme; Sigma-Aldrich) buffer (25 mM Tris pH 7.5, 10 mM EDTA, 585 mM sucrose, and 0.3 mg/ml sodium taurocholate) at room temperature and hybridized overnight at 46 °C in hybridization chambers with the oligonucleotide probe (final concentration of 5 ng/μl in a solution of 30 percent formamide, 0.9 M sodium chloride, 20 mM Tris pH 7.5, and 0.01% sodium dodecyl sulfate). Tissue sections were washed for 20 minutes at 48 °C in washing buffer (0.9 M NaCl, 20 mM Tris pH 7.2, 0.1% SDS, 20% Formamide, and 10% Dextran Sulfate) and once in distilled water for 10 seconds. The slide was stained with DAPI for 2 min and dried at RT, mounted with 50% glycerol. *C. perfringens* in intestinal tissue was visualized using a Nikon TS2 fluorescent microscopy and representative images of each treatment were presented in the Results section.

### Bile acid *C. perfringens* inhibition assay

*C. perfringens* in Tryptic Soy Broth (TSB) supplemented with 0.5% sodium thioglycollate, with added TCA (0.2 mM, final concentration), CA (0.2 mM) or DCA (0, 0.01, 0.05, 0.1, or 0.2 mM) or LCA or UDCA (0, 0.2, or 1 mM) was cultured at 42 °C overnight (16 -18 hours) under anaerobic conditions. The bacterial growth tubes were imaged for inhibition (clear broth) or no inhibition (cloudy broth) using a camera and representative images of each treatment were presented in the Results section. The bacterial growth was the quantified by OD600 nm using a spectrophotometer (Nanodrop, Thermo Fisher).

### Statistical Analysis

For *in vitro* assay of bile acids against *C. perfringens* growth, data were first analyzed by One-way ANOVA for significant difference and then Bonferroni’s multiple comparison test using Prism 5.0 software. For other data, differences between treatments were analyzed pairwise using the nonparametric Mann–Whitney *U* test performed using Prism 5.0 software. The specific pairwise comparisons were showed in Results section and Figures. Sample (individual birds or cell wells) numbers of body weight, histology, and other assays were listed in respective subsections of Methods. Values are shown as mean of samples in the treatment ± standard error of the mean as indicated. Experiments were considered statistically significant if *P* values were < 0.05.

## Supplemental Figure Legend

**Supplemental Figure 1. DCA attenuates NE-induced productivity loss.** Cohorts of thirteen broiler chicks were fed different diets and infected as in Figure 2. Bird body weight gain was measured at 18 (13 birds/group), 23 (13 birds/group), and 26 (8-9 birds/group) days of age. Showed was daily periodic body weight gain. All graphs depict mean ± SEM. **, P<0.01; ***, P<0.001. Results are representative of 3 independent experiments.

**Supplemental Figure 2. COX inhibitor aspirin alleviates murine cytokine-induced inflammatory response in chicken splenocytes.** Splenocytes isolated from broiler chickens were stimulated with murine mINFγ (1μg/ml) or mTNFα (5ng/ml) for 2 hr in the presence of 1.2 mM aspirin. RNA was extracted, reverse-transcribed, and quantified using a Bio-Rad 384 PCR platform.(A) *Infγ*, *Litaf*, and *Mmp9* mRNA fold change normalized to *Gapdh* in the presence of mINFγ. (B) *Infγ*, *Litaf*, and *Mmp9* mRNA fold change normalized to *Gapdh* in the presence of mTNFα. All graphs depict mean ± SEM. *, P<0.05. Results are representative of 2 independent experiments.

**Supplemental Figure 3. Aspirin attenuates NE-induced productivity loss.** Cohorts of 13 broiler chicks were fed different diets and infected as in Figure 6. Bird body weight gain was measured at 18 (13 birds/group), 23 (13 birds/group), and 26 (8 birds/group) days of age. Showed was daily periodic body weight gain. All graphs depict mean ± SEM. NS, not significant; *, P<0.01; Results are representative of 2 independent experiments.

## References

1. Neu HC. 1992. The crisis in antibiotic resistance. Science 257:1064–1073.

2. McGann P, Snesrud E, Maybank R, Corey B, Ong AC, Clifford R, Hinkle M, Whitman T, Lesho E, Schaecher KE. 2016. Escherichia coli Harboring mcr-1 and blaCTX-M on a Novel IncF Plasmid: First report of mcr-1 in the USA. Antimicrobial agents and chemotherapy.

3. Casewell M, Friis C, Marco E, McMullin P, Phillips I. 2003. The European ban on growth-promoting antibiotics and emerging consequences for human and animal health. The Journal of antimicrobial chemotherapy 52:159–161.

4. Wade B, Keyburn A. 2015. The true cost of necrotic enteritis. Poultry World 31:16–17.

5. Kaldhusdal M, Benestad SL, Lovland A. 2016. Epidemiologic aspects of necrotic enteritis in broiler chickens - disease occurrence and production performance. Avian pathology: journal of the W.V.P.A 45:271–274.

6. Timbermont L, Haesebrouck F, Ducatelle R, Van Immerseel F. 2011. Necrotic enteritis in broilers: an updated review on the pathogenesis. Avian Pathol 40:341–347.

7. Olkowski AA, Wojnarowicz C, Chirino-Trejo M, Drew MD. 2006. Responses of broiler chickens orally challenged with Clostridium perfringens isolated from field cases of necrotic enteritis. Research in veterinary science 81:99–108.

8. Olkowski AA, Wojnarowicz C, Chirino-Trejo M, Laarveld B, Sawicki G. 2008. Sub-clinical necrotic enteritis in broiler chickens: novel etiological consideration based on ultra-structural and molecular changes in the intestinal tissue. Research in veterinary science 85:543–553.

9. Prescott JF, Parreira VR, Mehdizadeh Gohari I, Lepp D, Gong J. 2016. The pathogenesis of necrotic enteritis in chickens: what we know and what we need to know: a review. Avian Pathol 45:288–294.

10. Caricilli AM, Castoldi A, Camara NO. 2014. Intestinal barrier: A gentlemen’s agreement between microbiota and immunity. World J Gastrointest Pathophysiol 5:18–32.

11. Subramanian S, Huq S, Yatsunenko T, Haque R, Mahfuz M, Alam MA, Benezra A, DeStefano J, Meier MF, Muegge BD, Barratt MJ, VanArendonk LG, Zhang Q, Province MA, Petri Jr Wa, Ahmed T, Gordon JI. 2014. Persistent gut microbiota immaturity in malnourished Bangladeshi children. Nature.

12. Belkaid Y, Hand TW. 2014. Role of the microbiota in immunity and inflammation. Cell 157:121‐141.

13. Deshmukh HS, Liu Y, Menkiti OR, Mei J, Dai N, O’Leary CE, Oliver PM, Kolls JK, Weiser JN, Worthen GS. 2014. The microbiota regulates neutrophil homeostasis and host resistance to Escherichia coli K1 sepsis in neonatal mice. Nat Med.

14. Rantala M, Nurmi E. 1973. Prevention of the growth of Salmonella infantis in chicks by the flora of the alimentary tract of chickens. British poultry science 14:627–630.

15. Silverman MS, Davis I, Pillai DR. 2010. Success of self-administered home fecal transplantation for chronic Clostridium difficile infection. Clin Gastroenterol Hepatol 8:471–473.

16. Buffie CG, Bucci V, Stein RR, McKenney PT, Ling L, Gobourne A, No D, Liu H, Kinnebrew M, Viale A, Littmann E, van den Brink Mr, Jenq RR, Taur Y, Sander C, Cross JR, Toussaint NC, Xavier JB, Pamer EG. 2015. Precision microbiome reconstitution restores bile acid mediated resistance to Clostridium difficile. Nature 517:205–208.

17. Ridlon JM, Kang DJ, Hylemon PB. 2006. Bile salt biotransformations by human intestinal bacteria. Journal of lipid research 47:241–259.

18. Ma H, Patti ME. 2014. Bile acids, obesity, and the metabolic syndrome. Best practice & research. Clinical gastroenterology 28:573–583.

19. Bernstein C, Holubec H, Bhattacharyya AK, Nguyen H, Payne CM, Zaitlin B, Bernstein H. 2011. Carcinogenicity of deoxycholate, a secondary bile acid. Archives of toxicology 85:863–871.

20. Sun X, Winglee K, Gharaibeh RZ, Gauthier J, He Z, Tripathi P, Avram D, Bruner S, Fodor A, Jobin C. 2018. Microbiota-Derived Metabolic Factors Reduce Campylobacteriosis in Mice. Gastroenterology 154:1751-1763 e1752.

21. Rose ME, Long PL, Bradley JW. 1975. Immune responses to infections with coccidia in chickens: gut hypersensitivity. Parasitology 71:357–368.

22. Dalloul RA, Lillehoj HS. 2006. Poultry coccidiosis: recent advancements in control measures and vaccine development. Expert Rev Vaccines 5:143–163.

23. Paiva D, McElroy A. 2014. Necrotic enteritis: Applications for the poultry industry. The Journal of Applied Poultry Research 23:557–566.

24. Uzal FA, Freedman JC, Shrestha A, Theoret JR, Garcia J, Awad MM, Adams V, Moore RJ, Rood JI, McClane BA. 2014. Towards an understanding of the role of Clostridium perfringens toxins in human and animal disease. Future Microbiol 9:361–377.

25. Freedman JC, Shrestha A, McClane BA. 2016. Clostridium perfringens Enterotoxin: Action, Genetics, and Translational Applications. Toxins 8.

26. Biondi C, Ferretti ME, Pavan B, Lunghi L, Gravina B, Nicoloso MS, Vesce F, Baldassarre G. 2006. Prostaglandin E2 inhibits proliferation and migration of HTR-8/SVneo cells, a human trophoblast-derived cell line. Placenta 27:592–601.

27. Dey I, Lejeune M, Chadee K. 2006. Prostaglandin E2 receptor distribution and function in the gastrointestinal tract. Br J Pharmacol 149:611–623.

28. Lazarus M. 2006. The differential role of prostaglandin E2 receptors EP3 and EP4 in regulation of fever. Mol Nutr Food Res 50:451–455.

29. Bos CL, Richel DJ, Ritsema T, Peppelenbosch MP, Versteeg HH. 2004. Prostanoids and prostanoid receptors in signal transduction. Int J Biochem Cell Biol 36:1187–1205.

30. Lee Y, Rodriguez C, Dionne RA. 2005. The role of COX-2 in acute pain and the use of selective COX-2 inhibitors for acute pain relief. Curr Pharm Des 11:1737–1755.

31. Wallace JL. 2001. Prostaglandin biology in inflammatory bowel disease. Gastroenterol Clin North Am 30:971–980.

32. Tessner TG, Muhale F, Riehl TE, Anant S, Stenson WF. 2004. Prostaglandin E2 reduces radiation-induced epithelial apoptosis through a mechanism involving AKT activation and bax translocation. J Clin Invest 114:1676–1685.

33. Martin-Venegas R, Roig-Perez S, Ferrer R, Moreno JJ. 2006. Arachidonic acid cascade and epithelial barrier function during Caco-2 cell differentiation. Journal of lipid research 47:1416–1423.

34. Zamuner SR, Warrier N, Buret AG, MacNaughton WK, Wallace JL. 2003. Cyclooxygenase 2 mediates post-inflammatory colonic secretory and barrier dysfunction. Gut 52:1714–1720.

35. Hatazawa R, Ohno R, Tanigami M, Tanaka A, Takeuchi K. 2006. Roles of endogenous prostaglandins and cyclooxygenase isozymes in healing of indomethacin-induced small intestinal lesions in rats. J Pharmacol Exp Ther 318:691–699.

36. Pavlidis P, Bjarnason I. 2015. Aspirin Induced Adverse Effects on the Small and Large Intestine. Curr Pharm Des 21:5089–5093.

37. Royer C, Lu X. 2011. Epithelial cell polarity: a major gatekeeper against cancer? Cell death and differentiation 18:1470–1477.

38. Xin ZS. 1998. An Atlas of Histology. Chapter 10. Digestive System, p. 188-270. Springer.

39. Delgado ME, Grabinger T, Brunner T. 2016. Cell death at the intestinal epithelial front line. The FEBS journal 283:2701–2719.

40. Grant TD, Specian RD. 2001. Epithelial cell dynamics in rabbit cecum and proximal colon P 1. The Anatomical record 264:427–437.

41. Nielsen OH, Rask-Madsen J. 1996. Mediators of inflammation in chronic inflammatory bowel disease. Scandinavian journal of gastroenterology. Supplement 216:149–159.

42. Sun X, Liu B, Sartor RB, Jobin C. 2013. Phosphatidylinositol 3-kinase-gamma signaling promotes Campylobacter jejuni-induced colitis through neutrophil recruitment in mice. J Immunol 190:357–365.

43. Sun X, Threadgill D, Jobin C. 2012. Campylobacter jejuni induces colitis through activation of mammalian target of rapamycin signaling. Gastroenterology 142:86-95 e85.

44. Caly DL, D’Inca R, Auclair E, Drider D. 2015. Alternatives to Antibiotics to Prevent Necrotic Enteritis in Broiler Chickens: A Microbiologist’s Perspective. Front Microbiol 6:1336.

45. Chapman HD, Barta JR, Blake D, Gruber A, Jenkins M, Smith NC, Suo X, Tomley FM. 2013. A selective review of advances in coccidiosis research. Advances in parasitology 83:93–171.

46. Abt MC, McKenney PT, Pamer EG. 2016. Clostridium difficile colitis: pathogenesis and host defence. Nature reviews. Microbiology 14:609–620.

47. Sorg JA, Sonenshein AL. 2008. Bile salts and glycine as cogerminants for Clostridium difficile spores. J Bacteriol 190:2505–2512.

48. Long JR, Pettit JR, Barnum DA. 1974. Necrotic enteritis in broiler chickens. II. Pathology and proposed pathogenesis. Canadian journal of comparative medicine: Revue canadienne de medecine comparee 38:467–474.

49. Al-Sheikhly F, Truscott RB. 1977. The pathology of necrotic enteritis of chickens following infusion of broth cultures of Clostridium perfringens into the duodenum. Avian diseases 21:230–240.

50. Sherman MA, Kalman D. 2004. Initiation and resolution of mucosal inflammation. Immunologic Research 29:241.

51. Winter SE, Thiennimitr P, Winter MG, Butler BP, Huseby DL, Crawford RW, Russell JM, Bevins CL, Adams LG, Tsolis RM, Roth JR, Baumler AJ. 2010. Gut inflammation provides a respiratory electron acceptor for Salmonella. Nature 467:426–429.

52. Hassan JO, Curtiss R, 3rd. 1994. Virulent Salmonella typhimurium-induced lymphocyte depletion and immunosuppression in chickens. Infect Immun 62:2027–2036.

53. Navarro MA, McClane BA, Uzal FA. 2018. Mechanisms of Action and Cell Death Associated with Clostridium perfringens Toxins. Toxins (Basel) 10.

54. Keyburn AL, Boyce JD, Vaz P, Bannam TL, Ford ME, Parker D, Di Rubbo A, Rood JI, Moore RJ. 2008. NetB, a new toxin that is associated with avian necrotic enteritis caused by Clostridium perfringens. PLoS Pathog 4:e26.

55. Martin TG, Smyth JA. 2009. Prevalence of netB among some clinical isolates of Clostridium perfringens from animals in the United States. Vet Microbiol 136:202–205.

56. Paiva DM, Walk CL, McElroy AP. 2013. Influence of dietary calcium level, calcium source, and phytase on bird performance and mineral digestibility during a natural necrotic enteritis episode. Poult Sci 92:3125–3133.

57. Galarza-Seeber R, Latorre JD, Bielke LR, Kuttappan VA, Wolfenden AD, Hernandez-Velasco X, Merino-Guzman R, Vicente JL, Donoghue A, Cross D, Hargis BM, Tellez G. 2016. Leaky Gut and Mycotoxins: Aflatoxin B1 Does Not Increase Gut Permeability in Broiler Chickens. Front Vet Sci 3:10.

58. Basak SC, Lee S, Barta JR, Fernando MA. 2006. Differential display analysis of gene expression in two immunologically distinct strains of Eimeria maxima. Parasitol Res 99:28–36.

59. Long PL, Millard BJ, Joyner LP, Norton CC. 1976. A guide to laboratory techniques used in the study and diagnosis of avian coccidiosis. Folia Vet Lat 6:201–217.

60. Shivaramaiah S, Wolfenden RE, Barta JR, Morgan MJ, Wolfenden AD, Hargis BM, Tellez G. 2011. The role of an early Salmonella Typhimurium infection as a predisposing factor for necrotic enteritis in a laboratory challenge model. Avian diseases 55:319–323.

61. McReynolds JL, Byrd JA, Anderson RC, Moore RW, Edrington TS, Genovese KJ, Poole TL, Kubena LF, Nisbet DJ. 2004. Evaluation of immunosuppressants and dietary mechanisms in an experimental disease model for necrotic enteritis. Poult Sci 83:1948–1952.

62. Heikinheimo A, Korkeala H. 2005. Multiplex PCR assay for toxinotyping Clostridium perfringens isolates obtained from Finnish broiler chickens. Letters in applied microbiology 40:407–411.

63. Tanaka M, Riddell RH. 1990. The pathological diagnosis and differential diagnosis of Crohn’s disease. Hepato-gastroenterology 37:18–31.

64. Odze R. 2003. Diagnostic problems and advances in inflammatory bowel disease. Modern pathology: an official journal of the United States and Canadian Academy of Pathology, Inc 16:347–358.

65. Sun X, Jobin C. 2014. Nucleotide-binding oligomerization domain-containing protein 2 controls host response to Campylobacter jejuni in Il10-/-mice. J Infect Dis 210:1145–1154.

66. Yuan JS, Reed A, Chen F, Stewart CN, Jr. 2006. Statistical analysis of real-time PCR data. BMC Bioinformatics 7:85.

67. Rueden CT, Schindelin J, Hiner MC, DeZonia BE, Walter AE, Arena ET, Eliceiri KW. 2017. ImageJ2: ImageJ for the next generation of scientific image data. BMC Bioinformatics 18:529.

68. Sangild PT, Siggers RH, Schmidt M, Elnif J, Bjornvad CR, Thymann T, Grondahl ML, Hansen AK, Jensen SK, Boye M, Moelbak L, Buddington RK, Westrom BR, Holst JJ, Burrin DG. 2006. Diet-and colonization-dependent intestinal dysfunction predisposes to necrotizing enterocolitis in preterm pigs. Gastroenterology 130:1776–1792.

